# NFYA regulates spermatogenesis through multifaceted chromatin interactions

**DOI:** 10.64898/2026.07.29.741030

**Authors:** Boyan Wang, Shan Jiang, Qianying Yang, Yi Zhang

## Abstract

During spermatogenesis, germ cells undergo extensive chromatin remodeling to promote highly diverse and stage-specific transcriptional programs that drive differentiation, meiosis, and spermiogenesis. However, conventional loss-of-function approaches do not allow systematic interrogation of gene function across spermatogenesis *in vivo*. Here, we introduce a protein-degron strategy coupled with intra-seminiferous tubule injection (dTAG-IS) to enable rapid, simultaneous, and stage-resolved functional investigation of NFYA during spermatogenesis. Acute NFYA-depletion induces apoptosis in differentiating spermatogonia (diff-Spg), pachytene spermatocytes (pSpc), and round spermatids (rStd), revealing an essential requirement across spermatogenesis. Mechanistically, NFYA exhibits stage-specific chromatin-binding patterns and directly regulates distinct transcriptional programs, including cell cycle progression and histone assembly in diff-Spg; chromosome organization and DNA damage response in pSpc; and antioxidative stress programs in rStd. Together, our study underscores dTAG-IS as a versatile strategy for *in vivo* loss-of-function studies during spermatogenesis and establishes NFYA as a multifaceted regulator that drives stage-specific transcriptional programs to ensure spermatogenesis progression.

## Introduction

Mammalian spermatogenesis is a tightly orchestrated process during which spermatogonial stem cells generate mature spermatozoa (Murat *et al*, 2022). In mice, spermatogenesis starts with the differentiation of spermatogonial stem cells to spermatogonia, which further give rise to pre-leptotene (preL) spermatocytes (Spc) that enter meiosis (Oakberg, 1956b). Meiosis is the most complex and critical phase of spermatogenesis, featuring an extremely prolonged prophase of meiosis I that spans four stages, including leptotene (L), zygotene (Z), pachytene (P) and diplotene (D) (Oakberg, 1956b; Soumillon *et al*, 2013) and lasts for two weeks in males. Completion of meiosis produces haploid round spermatids (rStd), which then undergo spermiogenesis to form mature spermatozoa (Oakberg, 1956a, b).

The differentiation process from Spg to mature spermatozoa is accompanied by profound chromatin remodeling, epigenetic programming, and dynamic transcriptional changes, which take roughly 35 days in mice (Ernst *et al*, 2019; Lin *et al*, 2025; Maezawa *et al*, 2018; Oakberg, 1956b). Thus, multiple germ-cell stages coexist within each seminiferous tubule in the adult testis. This cellular heterogeneity makes the study of individual sub-stages of spermatogenesis difficult using conventional Cre/loxp-based loss-of-function approaches because: 1) Cre/loxp system is not always efficient due to erratic recombination efficiency and the persistence of pre-existing protein obscures temporal resolution (Bao *et al*, 2013; Luo *et al*, 2020); 2) unexpected transient expression of Cre during early development may confound phenotypic interpretation (Luo *et al*., 2020; Song & Palmiter, 2018); 3) long-term gene ablation frequently leads to secondary effects that mask primary mechanisms; 4) systematic investigation across distinct spermatogenic stages requires multiple stage-specific Cre lines (Dai *et al*, 2024). To overcome these challenges, we took advantage of degron tag (dTAG)-mediated rapid protein degradation system (Nabet *et al*, 2018), which has been shown to be “non-toxic, rapid, and efficient” in embryos (Abuhashem *et al*, 2022; Zhou *et al*, 2025), and developed an *in vivo* loss-of-function dTAG system by coupling it with intra-seminiferous tubule injection (dTAG-IS). This strategy enables rapid, simultaneous, and stage-resolved functional investigation across spermatogenesis.

Previous studies focusing on the open chromatin dynamics during spermatogenesis have revealed a significant enrichment of NFYA motif in Spg and Spc (Guo *et al*, 2017; Huang *et al*, 2023; Maezawa *et al*., 2018; Wu *et al*, 2022), suggesting a potential role for NFYA during spermatogenesis. However, it remains unknown whether and how NFYA functions across the different stages of spermatogenesis. NFYA is the sequence-specific DNA binding subunit of the NFY/CCAAT complex with an evolutionarily conserved function in chromatin accessibility establishment or maintenance (Nardini *et al*, 2013; Oldfield *et al*, 2014). *Nfya* knockout is embryonic lethal (Bhattacharya *et al*, 2003). Here, as a proof-of-concept, we leveraged our recently established NFYA-dTAG mouse line (Yang *et al*, 2026) and combined dTAG-IS with RNA-seq and low-input CUT&RUN assays to systematically investigate the *in-situ* functions of NFYA across spermatogenesis. Phenotypically, acute NFYA-depletion causes apoptosis in Spg, Spc, and rStd, revealing an essential role of NFYA throughout spermatogenesis. Mechanistically, NFYA directly regulates genes governing cell cycle progression and histone assembly in diff-Spg through promoter binding. In pSpc, NFYA activates genes involved in chromosome organization and DNA damage response through both promoter and distal binding. In rStd, NFYA promotes antioxidative stress programs through promoter binding, thereby preventing accumulation of excessive intracellular reactive oxygen species (ROS). Thus, NFYA exhibits stage-specific chromatin occupancy that drives distinct transcriptional programs to safeguard spermatogenesis. Collectively, our study not only highlights dTAG-IS as a powerful strategy for *in vivo* stage-resolved loss-of-function analysis during spermatogenesis but also establishes NFYA as a multifaceted regulator orchestrating transcriptional networks to safeguard spermatogenesis.

## Results

### Establishment of dTAG-IS method for functional analysis across spermatogenesis

To explore the potential role of NFYA during spermatogenesis, we first analyzed public RNA-seq datasets across spermatogenesis (Lin *et al*., 2025) (**Fig. 1A**). We found that its transcription increases during Spg differentiation, declines in preL Spc before meiosis entry, and then rises again during meiosis and persists into rStd (**Fig. 1A**). Such expression dynamics is consistent with a potential role in spermatogenesis. Immunostaining showed that NFYA protein levels increase during meiosis, remains in the nucleus from Spg to diplotene Spc, is present in both the cytoplasm and nucleus at metaphase, becomes predominantly nuclear again in rStd, and then becomes undetectable after Step 7-8 (**Fig. 1B, upper panel**). To deplete NFYA in testis, we used our recently established NFYA-dTAG mouse model (Yang *et al*., 2026) and delivered dTAG^V^-1 by intraperitoneal (i.p.) injection for 6 or 12 h (**Fig. EV1A**). This strategy has been shown to be efficient in inducing targeted protein degradation in multiple tissues, including embryos, liver, intestine, heart and lung (Abuhashem *et al*., 2022). However, immunostaining revealed that NFYA was not depleted in the testis (**Fig. EV1B**), likely owing to the blood-testis barrier (Mruk & Cheng, 2015). To bypass this barrier, we directly delivered dTAG^V^-1 by intra-seminiferous tubule (IS) injection (**Fig. 1C**), a method previously used for efficient delivery of plasmids, adenoviruses, and donor germ cells (Hooley *et al*, 2008; Ogawa *et al*, 1997; Pramod & Mitra, 2018). Immunostaining revealed that NFYA was completely depleted 24h following IS injection of 20 μM dTAG^V^-1 (**Fig. EV1C**). To investigate the temporal dynamics of NFYA-depletion, we performed a time-course analysis and found that NFYA levels were largely reduced within 6h, completely depleted at 24h, and recovered by 48h after one-time dTAG^V^-1 treatment (**Fig. 1D-E**). A detailed quantification revealed that NFYA was efficiently depleted across all germ-cell types following IS injection of 20 μM dTAG^V^-1 for 24h (**Fig. 1B-E**), without significantly altering germ-cell numbers at each stage (**Fig. EV1D-E**). These results demonstrate that we have successfully established the dTAG-IS approach for analyzing stage-resolved function of NFYA during spermatogenesis.

**Figure 1.**
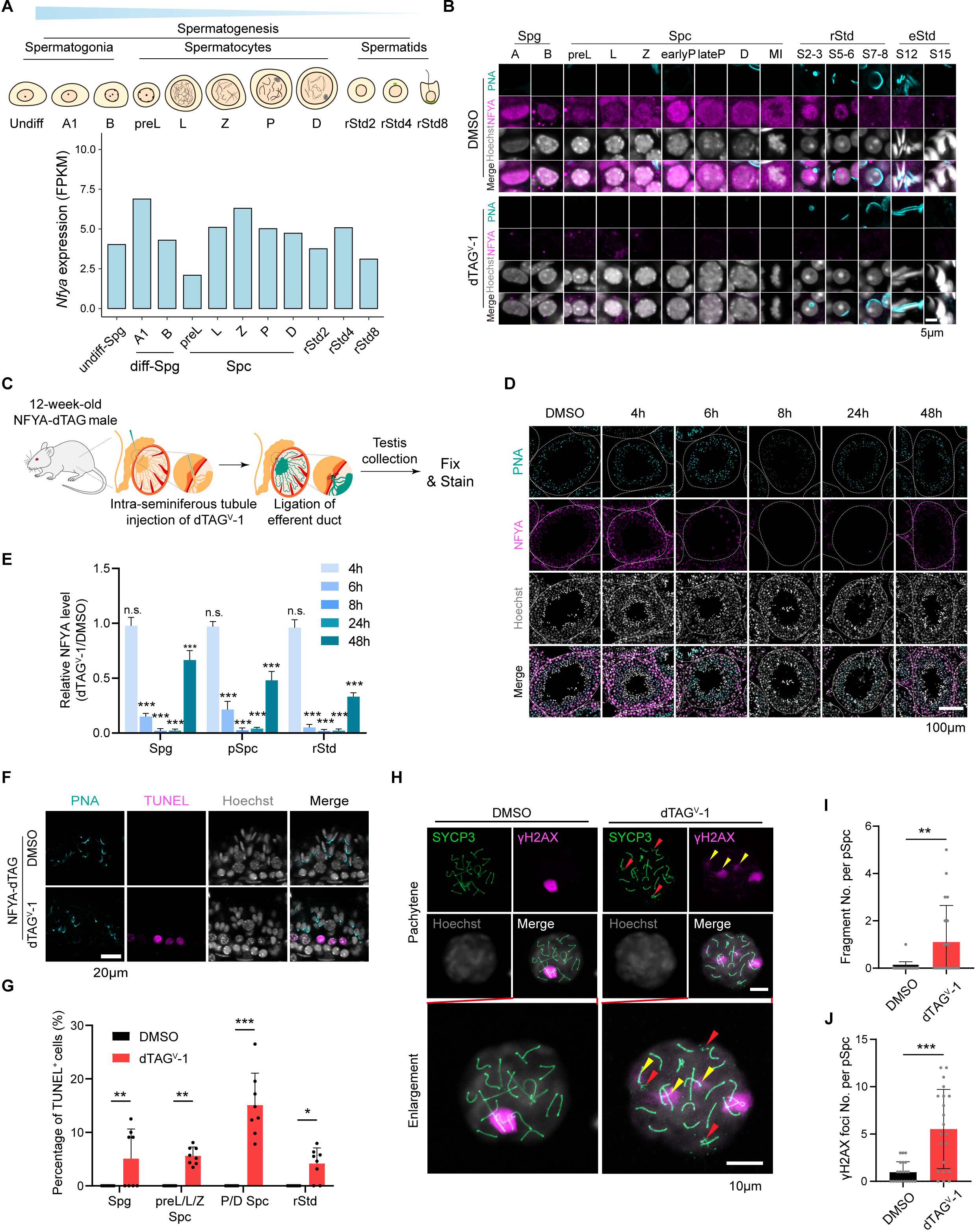
Acute NFYA degradation causes apoptosis during spermatogenesis. **(A)** Schematic representation of the stages of mouse spermatogenesis, including Spg, Spc, and rStd (Top). Bar graph showing *Nfya* mRNA expression levels across different stages of spermatogenesis based on RNA-seq data from GSE242515 (Bottom). **(B)** Immunofluorescence (IF) staining of HA-tagged NFYA (magenta) and PNA (cyan) in NFYA-dTAG testis sections treated with DMSO or dTAG^V^-1. Scale bar, 5 μm. **(C)** Experimental workflow for acute NFYA protein degradation *in vivo* via intra-seminiferous tubule (IS) injection. dTAG^V^-1 or DMSO was injected into the seminiferous tubules of 12-week-old NFYA-dTAG mice, followed by efferent duct ligation and testis collection at the indicated times. **(D)** IF images of HA-tagged NFYA (magenta) and PNA (cyan) in testis sections from NFYA-dTAG mice following DMSO or dTAGV-1 treatment for 4, 6, 8, 24, or 48 h. Dashed lines outline the seminiferous tubules, and other areas are the mesenchyme of testis. Scale bar, 100 μm. **(E)** Quantification of NFYA protein degradation kinetics. Mean fluorescence intensity of HA-NFYA is shown relative to DMSO at 4h, 6h, 8h, 24h, and 48h post-injection in Spg, pSpc, and rStd. **(F)** TUNEL assay (magenta) on testis sections 24h post-treatment with DMSO or dTAG^V^-1. Scale bar, 20 μm. **(G)** Quantification of the percentage of TUNEL^+^ cells in different spermatogenesis stages. **(H)** Immunostaining of SYCP3 (green) and γH2AX (magenta) with spermatocyte spreads from DMSO and dTAG^V^-1 treated testes. Red arrowheads indicate chromosome fragments; yellow arrowheads indicate abnormal γH2AX foci on autosomes. Scale bars, 10 μm. **(I-J)** Quantification of chromosome fragments (I) and abnormal γH2AX foci (J) per pachytene Spc. \**p* <0.05, ** *p* <0.01, *** *p* <0.001, n.s. *p* >0.05, *p* values were achieved using Student’s *t* test.

### Acute loss of NFYA causes apoptosis during spermatogenesis

Since spermatogenesis is a relatively long process, short-term protein depletion is unlikely to cause morphological or developmental phenotypes, including developmental arrest. We thus focused on apoptosis, a rapid and sensitive readout commonly triggered by failure of quality-control checkpoints that monitor cell-cycle progression, DNA damage repair, DNA integrity, synapsis and spindle assembly throughout spermatogenesis (Fang *et al*, 2021; Gartner *et al*, 2000; Odorisio *et al*, 1998; Sassone-Corsi, 1997). Thus, we performed TUNEL staining and found a significant increase in apoptosis of NFYA-depleted Spg, Spc, and rStd, especially in pachytene Spc (pSpc) (**Fig. 1F-G**), suggesting an essential requirement for NFYA in these cell types. Notably, IS injection of dTAG^V^-1 in WT mice did not increase apoptosis in germ cells (**Fig. EV1F**), excluding potential side-effects from dTAG^V^-1 injection itself. Intriguingly, we found that apoptosis increased more in P/D Spc upon NFYA-depletion (**Fig. 1G**). We therefore investigated DNA integrity in Spc and observed a significant increase in abnormal γH2AX foci (yellow arrowheads) and chromosome fragments (red arrowheads) in NFYA-depleted pSpc compared with the DMSO group (**Fig. 1H-J**). In contrast, no significant changes were observed in L, Z, and D Spc (**Fig. EV1G**), suggesting that pSpc are particularly vulnerable to NFYA-depletion. Taken together, these results indicate that NFYA-depletion triggers widespread apoptosis across spermatogenesis, with pSpc being especially sensitive.

### NFYA regulates genes involved in chromosome organization and DNA damage response

To understand how NFYA-depletion affects pSpc, we analyzed the transcriptional effect of NFYA degradation by RNA-seq. To this end, we performed dTAG-IS to deplete NFYA for 24h and isolated pSpc by Fluorescence-activated cell sorting (FACS) (**Fig. EV2A**). After confirming the high purity of sorted cells by immunostaining (**Fig. EV2B**), we performed RNA-seq (**Fig. EV2C**). Comparative analysis (Fold Change > 2 & FDR < 0.05) identified 372 downregulated and 390 upregulated genes in pSpc upon NFYA-depletion, whereas repeat elements showed little change (**Fig. 2A and Dataset EV1**). Gene Ontology (GO) analysis revealed that downregulated genes are enriched for chromosome organization and upregulated genes are enriched for apoptotic process (**Fig. 2B**), consistent with the observed chromosome fragmentation and apoptosis phenotypes in pSpc. Additionally, gene set enrichment analysis (GSEA) revealed that DNA damage recognition and RNA polymerase III (POLIII) transcription were significantly downregulated in NFYA-depleted pSpc (**Fig. 2C and Fig. EV2D**), consistent with observed abnormal DNA damage in pSpc. Furthermore, examination of mouse genome information (MGI) phenotypic annotation of downregulated genes revealed that NFYA-depletion affected many genes, including *Dmrtc2*, *Cdkn2d*, and *Cdca8* (**Fig. 2D**), whose knockouts or mutations exhibit abnormal spermatogenesis, infertility, and apoptotic phenotypes, supporting a critical role of NFYA in pSpc.

**Figure 2.**
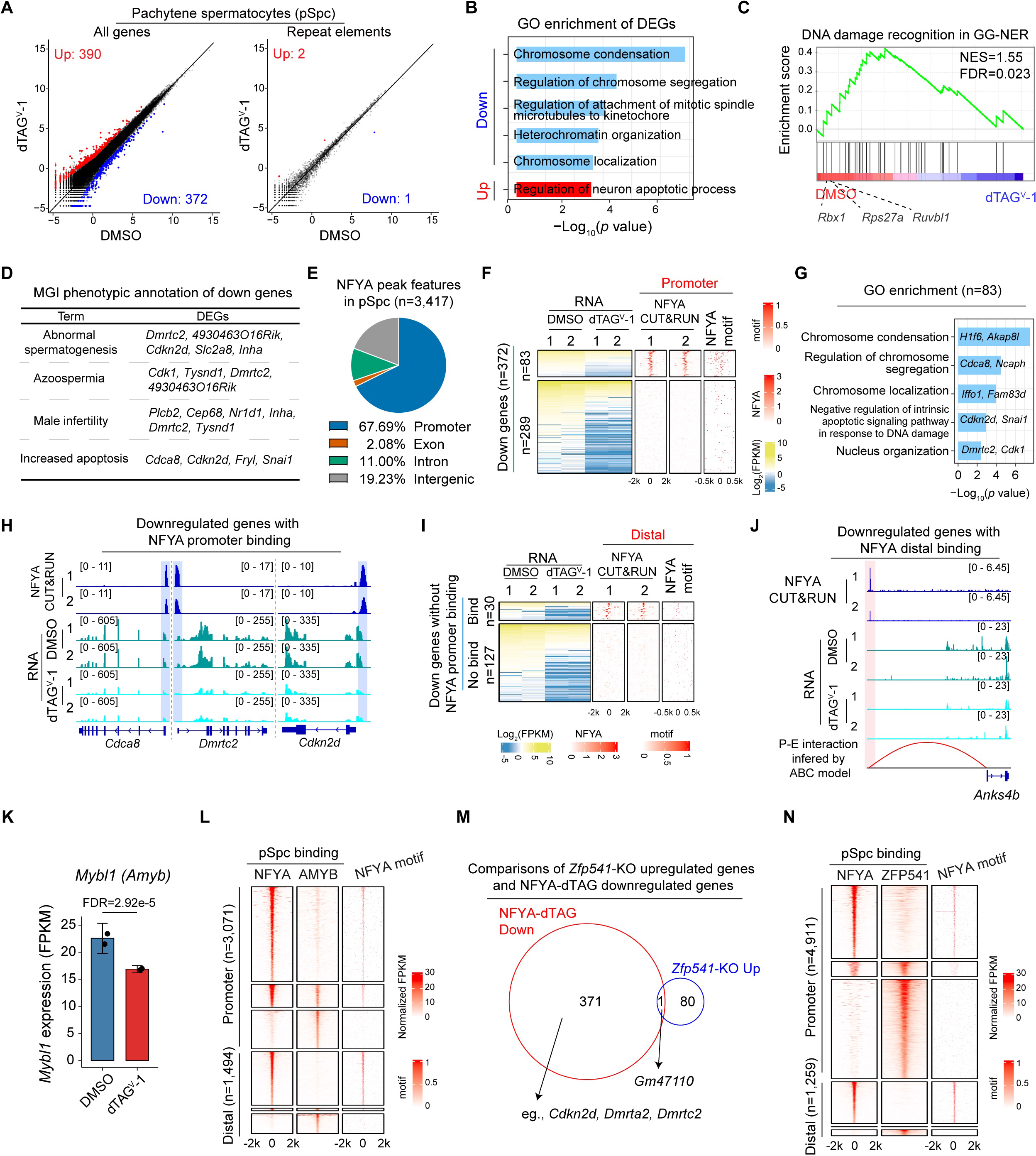
NFYA-depletion results in dysregulation of genes involved in chromosome organization and DNA damage response in pachytene spermatocytes. **(A)** Scatter plots comparing gene expression and repeat element expression profiles in pSpc following IS injection of DMSO or dTAG^V^-1. The x-and y-axes indicate Log_2_CPM values from RNA-seq. Differentially expressed genes were defined as those with fold change≥2 and FDR<0.05. **(B)** Representative GO terms enriched among downregulated (n=372) and upregulated genes (n=390) following NFYA depletion. Example genes for the highlighted terms are shown. **(C)** GSEA showing that genes involved in DNA damage recognition during global genome nucleotide excision repair (GG-NER) are significantly downregulated in dTAG^V^-1-treated pSpc. **(D)** MGI phenotype annotation of downregulated genes (n=372) in pSpc following dTAG^V^-1 IS injection. **(E)** Genomic distribution of NFYA-binding peaks (n=3,417) identified by NFYA CUT&RUN in pSpc. **(F)** Heatmaps showing differentially expressed genes in pSpc following NFYA depletion, together with NFYA binding and NFYA motif occurrence around the transcription start sites (TSSs) of the corresponding genes. Downregulated genes are separated into two groups based on the presence or absence of direct NFYA promoter binding. **(G)** GO terms enriched among NFYA-bound downregulated genes (n=83), with representative examples shown. **(H)** Genome browser views of representative NFYA target genes that are downregulated upon NFYA depletion, showing NFYA CUT&RUN and RNA-seq signals in pSpc treated with DMSO or dTAG^V^-1. **(I)** Heatmaps of downregulated genes lacking NFYA promoter binding, together with NFYA CUT&RUN and NFYA motif occurrence at distal regulatory regions. Genes are separated into two groups based on the presence or absence of distal NFYA binding. **(J)** Genome browser view of a representative distally regulated NFYA target gene in pSpc, showing NFYA CUT&RUN and RNA-seq signals, as well as the predicted promoter-enhancer interaction inferred by the ABC model. **(K)** Bar plot showing *Amyb* (*Mybl1*) expression levels in DMSO and dTAG^V^-1 treated pSpc. **(L)** Heatmaps showing NFYA and AMYB binding at promoter-proximal (n=3,071) and distal (n=1,494) regions in pSpc, together with NFYA motif occurrence. **(M)** Venn diagram showing the overlap between genes downregulated in NFYA-depleted pSpc and genes upregulated in *Zfp541*-KO pSpc. **(N)** Heatmaps showing NFYA and ZFP541 binding at proximal (n=4,911) and distal (n=1,259) regions of promoters in pSpc, together with NFYA motif occurrence.

To identify the differentially expressed genes (DEGs) directly affected by NFYA-depletion, we performed CUT&RUN experiments in pSpc and found 67.69% NFYA peaks are mapped to gene promoter regions (**Fig. 2E and Fig. EV2E**). Importantly, the NFYA binding peaks are highly enriched for NFYA binding motif (**Fig. EV2F-G**), demonstrating the specificity of the binding peaks. Furthermore, integrative analysis revealed that 83 out of 372 downregulated gene promoters are occupied by NFYA (**Fig. 2F and Dataset EV1**). Since low-input CUT&RUN can inherently miss peaks leading to underestimation of binding sites, we performed motif analysis on the 289 downregulated genes without detectable CUT&RUN signal and found that 79.9% (231/289) contained at least one NFYA motif in their promoter regions (**Fig. EV2H**), suggesting a direct role of NFYA in regulating these genes. GO analysis of these 83 direct targets, such as *Cdca8*, *Dmrtc2*, and *Cdkn2d*, revealed enrichment in terms related to chromosome organization and DNA damage response (**Fig. 2G-H**), which are essential for pSpc regulation. These data collectively suggest that NFYA directly regulates genes involved in chromosome organization and DNA damage response in pSpc.

Since 32.31% NFYA peaks are located at distal regions (putative enhancers) (**Fig. 2E and Fig. EV2F**), we next asked whether those distal binding events are also correlated with gene regulation. Cumulative distribution analysis revealed that distal NFYA-binding sites were located closer to downregulated genes, particularly those with NFYA promoter binding (**Fig. EV2I**), suggesting that NFYA may also regulate these genes through distal binding. To further characterize NFYA distal binding, we utilized the activity-by-contact (ABC) model (Fulco *et al*, 2019) to predict potential NFYA-mediated promoter-enhancer (P-E) interactions (**Fig. EV2J**). This analysis indicated that many NFYA-bound enhancers are linked to downregulated genes without detectable promoter NFYA CUT&RUN signal, including *Anks4b* (**Fig. 2I-J**). Collectively, our integrated analysis supports that NFYA directly regulates genes in pSpc by both promoter and distal binding.

### NFYA regulates transcriptional programs in pSpc independent of AMYB and ZFP541

Previous studies have identified AMYB (also known as MYBL1) (Bolcun-Filas *et al*, 2011; Li *et al*, 2013; Sassone-Corsi, 1997) as a master activator of pSpc, whereas ZFP541 (Horisawa-Takada *et al*, 2021; Xu *et al*, 2022) functions as a repressor of pachytene transcriptional programs. We therefore asked whether NFYA is functionally linked to AMYB or ZFP541. Acute NFYA degradation significantly decreased *Amyb* expression in pSpc (**Fig. 2K**), suggesting that NFYA may act upstream of *Amyb*. However, because NFYA-depletion only reduces 26% of *Amyb* expression and *Amyb* heterozygous knockout mice do not show spermatogenesis defects (Bolcun-Filas *et al*., 2011; Li *et al*., 2013), we propose that the NFYA-regulated transcriptional programs, such as chromosome organization and DNA damage response, are largely independent of AMYB. Since no public RNA-seq data from *Amyb*-KO pSpc is available, we compared their binding profiles and found that NFYA and AMYB bind to different genomic sites (**Fig. 2L**). Similar analysis revealed that NFYA-depletion did not affect *Zfp541* expression in pSpc (**Fig. EV2K**). Additionally, an integrated analysis further showed that only 1 out of 372 NFYA-downregulated genes overlapped with those of ZFP541-repressed genes (**Fig. 2M**), suggesting that they function independently. Consistently, comparison of NFYA and ZFP541 binding profiles revealed distinct binding patterns (**Fig. 2N**). Collectively, these results support that NFYA regulates transcriptional programs in pSpc independent of AMYB and ZFP541.

### NFYA regulates cell cycle progression and histone genes in differentiating spermatogonia

To understand NFYA’s role in Spg, we depleted NFYA for 24h by dTAG-IS and isolated differentiating spermatogonia (diff-Spg) by FACS using cKIT staining (**Fig. EV3A-B**), which is widely used for diff-Spg isolation (Maezawa *et al*., 2018). RNA-seq (**Fig. EV3C**) revealed 329 downregulated and 203 upregulated genes in cKIT^+^ Spg upon NFYA-depletion, whereas repeat elements showed no changes (**Fig. 3A and Dataset EV2**). GO analysis revealed that downregulated genes are enriched for nucleosome assembly and upregulated genes are enriched for apoptotic process (**Fig. 3B**), consistent with observed phenotypes. Importantly, MGI phenotypic annotation of downregulated genes revealed that loss-function of these affected genes exhibit abnormal spermatogenesis, infertility, and apoptosis phenotypes (**Fig. 3C**), consistent with a critical role of NFYA in regulating cKIT^+^ Spg. Intriguingly, GSEA revealed significant downregulation of histone genes in NFYA-depleted cKIT^+^ Spg (**Fig. 3D**), consistent with dysregulation of nucleosome assembly identified in GO analysis (**Fig. 3B**). Although histone genes are essential for spermatogenesis (Fontaine *et al*, 2022; Kennedy *et al*, 1985; Teng *et al*, 2009; Zhang *et al*, 2016), little is known about how histone genes are regulated. To determine the role of NFYA in histone gene regulation, we first performed a systematic analysis of histone gene expression dynamics and identified 3 clusters (**Fig. EV3D and Dataset EV3**). We then focused on C1 which increases from undiff-Spg to diff-Spg. DEGs analysis revealed that 21 out of 58 C1 histone genes were downregulated upon NFYA-depletion in cKIT^+^ Spg, including 4 H1 subtypes, 5 H2A subtypes, 5 H2B subtypes, and 7 H3 subtypes (**Fig. 3E and Fig. EV3E-I**), suggesting that NFYA is a key regulator of histone gene upregulation during Spg differentiation.

**Figure 3.**
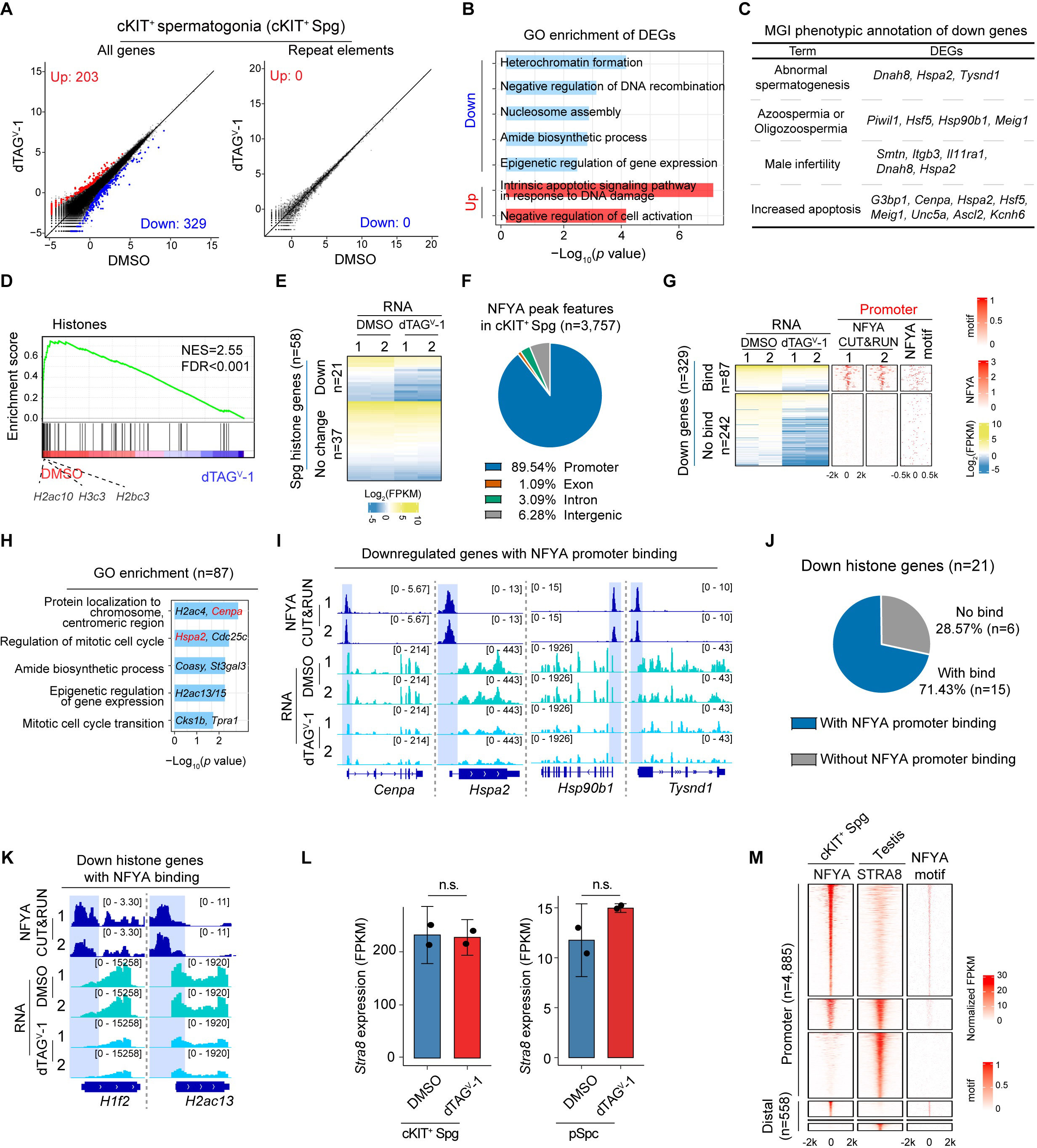
NFYA regulates cell cycle progression and histone gene upregulation in differentiating spermatogonia. **(A)** Scatter plots comparing gene expression and repeat element expression profiles in cKIT^+^ Spg following IS injection of DMSO or dTAG^V^-1. The x-and y-axes indicate Log_2_CPM values from RNA-seq. Differentially expressed genes were defined as those with fold change≥2 and FDR<0.05. **(B)** Representative GO terms enriched among downregulated (n=329) and upregulated genes (n=203) after dTAG^V^-1 treatment. **(C)** MGI phenotype annotation of the downregulated genes (n=329) following dTAG^V^-1 IS injection. **(D)** GSEA showing that histone genes are significantly downregulated in dTAG^V^-1-treated cKIT^+^ Spg. **(E)** Heatmap showing expression of histone genes in Spg (n=58) in cKIT^+^ Spg after NFYA depletion. **(F)** Genomic distribution of NFYA-binding peaks (n=3,757) identified by NFYA CUT&RUN in cKIT^+^ Spg. **(G)** Heatmaps showing differentially expressed genes in cKIT^+^ Spg following NFYA depletion, together with NFYA binding and NFYA motif occurrence around the TSSs of the corresponding genes. Downregulated genes are separated into two groups based on the presence or absence of direct NFYA promoter binding. **(H)** GO terms enriched among NFYA-bound downregulated genes (n=87), with representative examples shown. **(I)** Genome browser views of representative NFYA target genes that are downregulated upon NFYA depletion, showing NFYA CUT&RUN and RNA-seq signals in cKIT^+^ Spg treated with DMSO or dTAG^V^-1. **(J)** Pie chart showing the proportion of downregulated histone genes after NFYA depletion that are either bound or not bound by NFYA. **(K)** Genome browser views of representative downregulated histone genes with NFYA binding, showing NFYA CUT&RUN and RNA-seq signals at the H1f2 and H2ac13 loci in cKIT^+^ Spg treated with DMSO or dTAG^V^-1. **(L)** Bar plots showing *Stra8* expression levels in DMSO and dTAG^V^-1 treated cKIT^+^ Spg and pSpc. n.s., FDR >0.05. **(M)** Heatmaps showing NFYA, STRA8, and NFYA motif signals around promoter-proximal and distal regions in cKIT^+^ Spg and testis.

To identify NFYA’s direct targets in diff-Spg, we performed CUT&RUN experiments in cKIT^+^ Spg and found that 89.54% NFYA peaks are mapped to promoter regions (**Fig. 3F and Fig. EV3J**). Importantly, the presence of highly enriched NFYA motif in the identified NFYA binding peaks demonstrated the specificity of the NFYA bindings (**Fig. EV3K-L**). Furthermore, integrative analysis revealed that 87 out of 329 downregulated gene promoters are bound by NFYA (**Fig. 3G and Dataset EV2**). Since low-input CUT&RUN tends to underestimate binding events, we performed motif analysis of the 242 downregulated genes without detectable promoter NFYA CUT&RUN signal and identified 80.1% (194/242) of these genes contained at least one NFYA binding motif in their promoter regions (**Fig. EV3M**), suggesting that NFYA likely regulates these genes directly through promoter binding. GO analysis of the 87 downregulated direct targets, such as *Cenpa*, *H2ac4*, *Hspa2*, *Hsp90b1*, and *Tysnd1*, revealed enrichment in terms related to cell cycle progression (**Fig. 3H-I**), which is essential checkpoints (Fang *et al*., 2021; Sassone-Corsi, 1997) for Spg development. Importantly, we further identified 71.43% (15/21) downregulated histone genes were directly targeted by NFYA (**Fig. 3J-K and Fig. EV3N**), suggesting that NFYA is indeed a master regulator for histone gene expression during Spg differentiation. Together, these results suggest that NFYA directly regulates transcriptional programs, including cell cycle progression and histone gene upregulation, in diff-Spg.

Although STRA8 has previously been defined as a master regulator of meiosis initiation, recent studies revealed its important role in diff-Spg (Endo *et al*, 2015; Gewiss *et al*, 2021; Kojima *et al*, 2019). Consistently, *Stra8* expression is dramatically increased from undiff-Spg to diff-Spg (**Fig. EV3O**). We therefore asked whether NFYA is functionally linked to STRA8. We found that NFYA-depletion does not affect *Stra8* expression in cKIT^+^ Spg and pSpc (**Fig. 3L**). Furthermore, comparison of NFYA and STRA8 binding profiles revealed distinct binding patterns (**Fig. 3M**). Collectively, these results support that NFYA regulates transcriptional programs in diff-Spg independent of STRA8.

### NFYA directly regulates antioxidative stress programs in round spermatids

After establishing NFYA’s critical function in diff-Spg and pSpc, we next investigated its function in rStd. We used the same dTAG-IS approach to deplete NFYA for 24h and then isolated rStd by FACS (**Fig. EV4A-B**) for RNA-seq. We identified 140 downregulated and 126 upregulated genes in rStd in response to NFYA loss, while no obvious effect was identified for repeat elements (**Fig. 4A, Fig. EV4C, and Dataset EV4**). GO analysis revealed that downregulated genes are enriched for “response to endoplasmic reticulum (ER) stress” and upregulated genes are enriched for apoptotic process (**Fig. 4B**), consistent with observed phenotypes. Importantly, MGI phenotypic annotation revealed that loss-function of many downregulated genes, such as *Dmrtc2*, *Pih1d3*, and *Sesn2*, exhibit abnormal spermatogenesis, infertility, and apoptosis phenotypes (**Fig. 4C**), supporting a critical role of NFYA in rStd.

**Figure 4.**
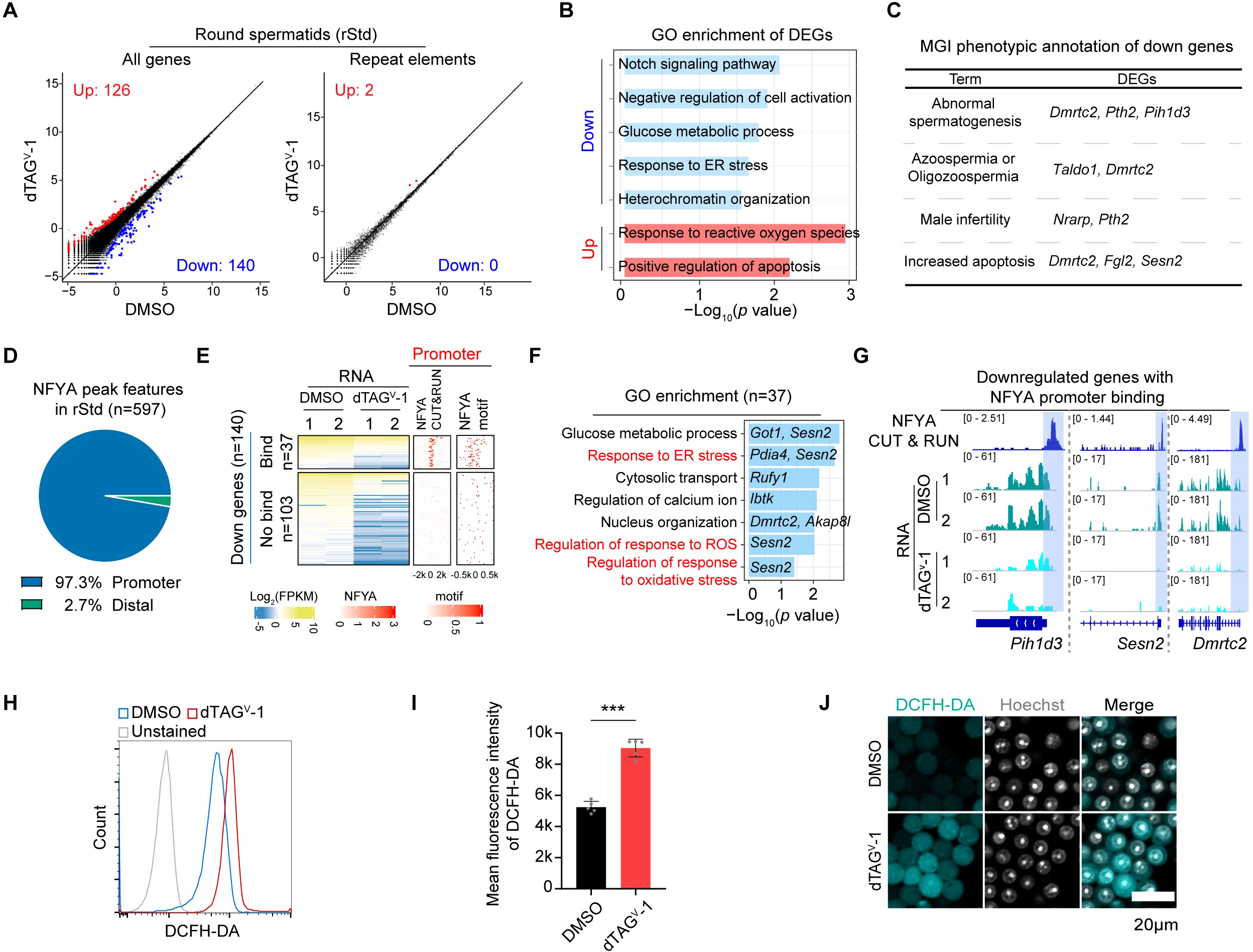
NFYA directly regulates genes involved in oxidative stress response in round spermatids. **(A)** Scatter plots comparing gene expression and repeat element expression profiles in rStd following IS injection of DMSO or dTAG^V^-1. The x-and y-axes indicate Log_2_CPM values from RNA-seq. Differentially expressed genes were defined as those with fold change≥2 and FDR<0.05. **(B)** Representative GO terms enriched among downregulated (n=140) and upregulated genes (n=126) following NFYA depletion. Example genes for the highlighted terms are shown. **(C)** MGI phenotype annotation of downregulated genes (n=140) in rStd following dTAG^V^-1 IS injection. **(D)** Genomic distribution of NFYA-binding peaks (n=597) identified by NFYA CUT&RUN in rStd. **(E)** Heatmaps showing downregulated genes in rStd following NFYA depletion, together with NFYA CUT&RUN signals and NFYA motif occurrence around the TSSs of the corresponding genes. Downregulated genes are separated into two groups based on the presence or absence of direct NFYA promoter binding. **(F)** GO terms enriched among NFYA-bound downregulated genes (n=37), with representative examples shown. **(G)** Genome browser views of representative NFYA target genes that are downregulated upon NFYA depletion, showing NFYA CUT&RUN and RNA-seq signals in rStd treated with DMSO or dTAG^V^-1. **(H)** Flow cytometric analysis of DCFH-DA fluorescence in rStd isolated from DMSO and dTAG^V^-1 treated testes. Unstained cells were used as a negative control. **(I)** Quantification of DCFH-DA mean fluorescence intensity in rStd after DMSO or dTAG^V^-1 treatment. *p* value is calculated using Student’s *t* test, \*\*\**p <*0.001. **(J)** Representative fluorescence images of sorted rStd stained with DCFH-DA and Hoechst after DMSO or dTAG^V^-1 treatment. Scale bar, 20 μm.

To identify genes directly affected by NFYA loss, we performed NFYA CUT&RUN in rStd (**Fig. EV4D**) and found that NFYA mainly occupies the promoter regions (**Fig. 4D**) that are highly enriched for NFYA binding motif (**Fig. EV4E-F**). Furthermore, integrative analysis revealed 37 out of the 140 downregulated genes are bound by NFYA (**Fig. 4E and Dataset EV4**), which is consistent with its motif enrichment in these promoters. Since low-input CUT&RUN tends to underestimate binding events, we performed motif analysis of the 103 downregulated genes without detectable promoter NFYA CUT&RUN signal and found 89.3% (92/103) of these genes contained at least one NFYA binding motif in their promoter regions (**Fig. EV4G**), supporting a likely direct role for NFYA-promoter binding in their regulation. GO analysis of the 37 downregulated direct targets, including *Sesn2* and *Dmrtc2*, revealed enrichment for pathways related to response to ER stress, ROS, and oxidative stress (**Fig. 4F-G and Fig. EV4H**). Notably, SESN2 (Sestrin2) is a highly conserved antioxidant protein that directly scavenges ROS and rectifies oxidative damage (Lu *et al*, 2023). Consistently, *Sesn2*-KO mice exhibit elevated oxidative stress and germ-cell loss (Chen *et al*, 2024). We therefore asked whether NFYA-depletion induces oxidative stress in rStd by measuring the ROS levels. We found a significant increase in the ROS levels in NFYA-depleted rStd compared with controls (**Fig. 4H-J**), suggesting that NFYA plays a critical role in maintaining oxidative homeostasis in rStd. Taken together, these data indicate that NFYA directly regulates antioxidative stress programs in rStd.

### NFYA regulates spermatogenesis by multifaceted chromatin binding

Having demonstrated that NFYA is a key transcription factor regulating multiple stages of spermatogenesis, we next sought to identify common and stage-specific roles of NFYA in diff-Spg, pSpc, and rStd. Comparison of downregulated genes (FDR < 0.05 & -Log_2_FC > 1) across the three stages revealed almost no overlap (**Fig. EV5A)**. Even when applying a more permissive threshold (FDR < 0.05 & -Log_2_FC > 0), only minimal overlap was detected (**Fig. EV5B)**, suggesting NFYA regulates distinct transcriptional programs at these stages. Next, we asked whether these differences reflect dynamic NFYA–chromatin interactions. Comparison of NFYA binding profiles identified 5 clusters (C1-C5), with 73.4% (3,668/4,996) of NFYA binding displaying stage-specific features (**Fig. 5A**). Notably, C1, C2, and C5 were predominantly located at promoter regions, whereas C3 and C4 were mainly located at distal regions (**Fig. 5B-C**), underscoring stage-specific NFYA chromatin-binding preferences. To further characterize their features, we performed integrative analyses of CUT&RUN and RNA-seq datasets to identify cluster-associated downregulated genes across spermatogenesis (**Fig. 5D-F and Dataset EV5**). For example, in cKIT^+^ Spg we identified 169 C1-related, 67 C2-related, and 215 C5-related downregulated genes (**Fig. 5D**). Consistently, comparative analysis revealed minimal overlap among NFYA-binding-related downregulated genes across these stages (**Fig. 5G**). Similar patterns were observed for C2-, C4-, and C5-related downregulated genes (**Fig. 5H and Fig. EV5C-D**). These findings suggest that multifaceted NFYA-chromatin interactions regulate distinct transcriptional programs at different stages of spermatogenesis. Consistent with this notion, GO analysis of stage-specific downregulated genes revealed enrichment for distinct biological processes, including “cell cycle progression” and “nucleosome assembly” in cKIT^+^ Spg, “chromosome segregation” and “DNA damage response” in pSpc, “response to ER stress” and “spermatid development” in rStd (**Fig. 5I**). Together, these results support stage-specific functions of NFYA binding across spermatogenesis.

**Figure 5.**
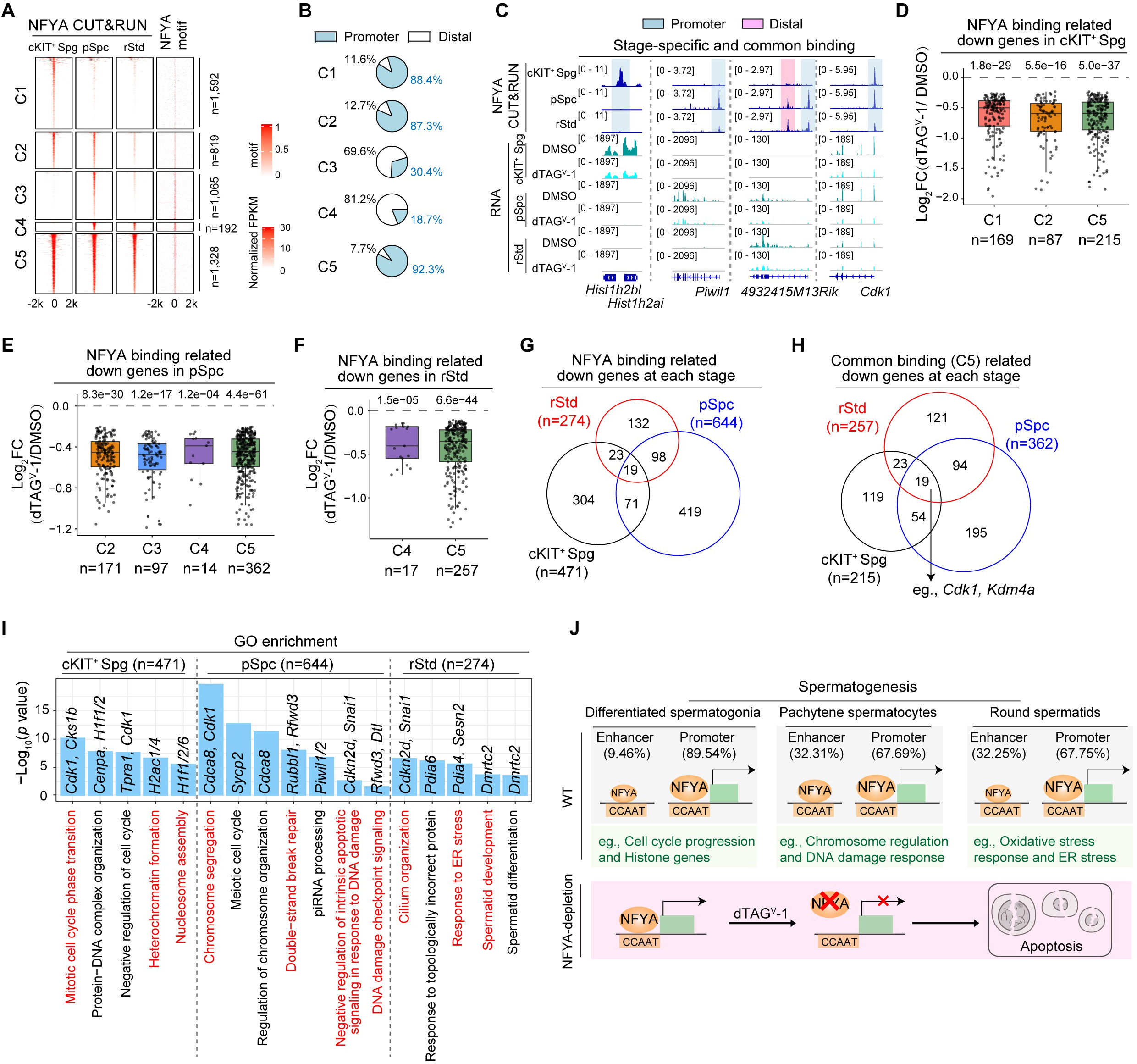
Multifaceted NFYA chromatin binding regulates spermatogenesis. **(A)** Heatmaps showing NFYA CUT&RUN signals across cKIT^+^ Spg, pSpc, and rStd, together with NFYA motif occurrence, in the five clusters of NFYA-binding regions (C1-C5) identified by comparative analysis across the three spermatogenesis stages. Numbers of genes in each cluster are indicated on the right. **(B)** Pie charts showing the genomic distribution of NFYA-binding regions in each gene cluster. **(C)** Genome browser views of representative stage-specific and common NFYA-binding regions, showing NFYA CUT&RUN and RNA-seq signals in cKIT^+^ Spg, pSpc, and rStd. Proximal or distal NFYA-binding regions are highlighted. **(D-F)** Box plots showing expression changes of NFYA-binding genes in cKIT^+^ Spg (D), pSpc (E), and rStd (F), grouped by NFYA-binding clusters. The number of genes in each group is indicated below. *p* values were calculated using the Wilcoxon test and are shown above each comparison. **(G)** Venn diagram showing the overlap of NFYA-binding-related downregulated genes across cKIT^+^ Spg, pSpc, and rStd. **(H)** Venn diagram showing the overlap of common binding (C5) related downregulated genes across cKIT^+^ Spg, pSpc, and rStd. Representative shared target genes are indicated. **(I)** Representative GO terms enriched among stage-specific NFYA-binding-related downregulated genes in cKIT^+^ Spg, pSpc, and rStd, with representative genes shown. **(J)** Model illustrating how NFYA exhibits multifaceted chromatin-binding patterns across spermatogenesis to regulate distinct stage-specific transcriptional programs, including cell cycle progression and histone gene expression in cKIT^+^ Spg, chromosome organization and DNA damage response in pSpc, and oxidative stress response in rStd.

Furthermore, motif enrichment analysis of these functional NFYA binding regions revealed, in addition to the NFYA motif, enrichment of TF binding motifs of some well-known stage-specific transcription factors (**Fig. EV5E-G**). These included TCF3, CTCFL, and ZBTB16 at cKIT^+^ Spg-associated NFYA-binding regions; TCFL5, MYBL1 (AMYB) and PKNOX1 at pSpc-associated NFYA-binding regions; RFX1/2 and SOX30 at rStd-associated NFYA-binding regions (**Fig. EV5E-G**). These findings suggest that these stage-specific TFs may help shape the multifaceted NFYA chromatin binding profiles during spermatogenesis. Future studies can test this possibility.

## Discussion

Testis exhibits one of the most complex and diverse transcriptomes among mammalian organs, even more complex than brain (Brawand *et al*, 2011; Soumillon *et al*., 2013). This complexity arises from extensive transcriptional reprogramming during spermatogenesis, which makes the study of stage-specific transcriptional regulation using conventional Cre/loxP-based loss-of-function technically challenging. Here we develop the dTAG-IS approach, which makes rapid, simultaneous, and stage-resolved functional investigation across different stages of spermatogenesis possible. As a proof-of-concept, we systematically investigated NFYA’s function across spermatogenesis. Phenotypically, acute NFYA-depletion causes widespread apoptosis across different stages of spermatogenesis, revealing its essential function. Mechanistically, NFYA exhibits multifaceted chromatin-binding patterns and directly regulates cell cycle progression and histone transcription in diff-Spg, chromosome organization and DNA damage response in pSpc, and antioxidative stress programs in rStd (**Fig. 5J**). Taken together, these findings highlight dTAG-IS as a versatile strategy for *in vivo* loss-of-function studies during spermatogenesis and establish NFYA as a multifaceted regulator that drives stage-specific transcriptional programs to ensure normal spermatogenesis progression.

Although previous motif enrichment analyses have suggested a potential role for NFYA in spermatogenesis (Guo *et al*., 2017; Huang *et al*., 2023; Maezawa *et al*., 2018; Wu *et al*., 2022), whether NFYA indeed functions during spermatogenesis and how NFYA functions remained unknown. Intriguingly, a recent preprint reported that scRNA-seq of testis from *Stra8*-Cre *Nfya*^fl/fl^ mouse only captured undiff-Spg (Säflund *et al*, 2026), suggesting a complete loss of diff-Spg and implicating NFYA in Spg differentiation. However, the mechanisms by which NFYA regulates diff-Spg were not addressed. Taking advantage of the dTAG-IS approach, we show that NFYA directly promotes histone gene expression and cell-cycle progression through promoter binding, and that acute NFYA depletion induces apoptosis in diff-Spg. Consistent with previous reports in other cell types (Cui *et al*, 2023; Dolfini *et al*, 2024), NFYA directly regulates cell cycle progression in diff-Spg. Although histone genes are essential for spermatogenesis (Fontaine *et al*., 2022; Kennedy *et al*., 1985; Teng *et al*., 2009; Zhang *et al*., 2016), the mechanisms governing their regulation during this process remain poorly understood. Here we reveal that NFYA regulates 21 out of 58 histone genes, demonstrating NFYA as a master regulator for histone gene upregulation during Spg differentiation. Notably, NFYA-depletion does not significantly affect histone gene expression in pSpc, suggesting a stage-specific role of NFYA in diff-Spg and indicating that other factors may compensate for histone gene regulation at later stages.

During mouse spermatogenesis, pachytene phase is one of the most complex and transcriptionally active stages, lasting for about 6 days (Goetz *et al*, 1984; Oakberg, 1956b) and characterized by massive chromosome organization, crossover formation, XY-body formation, meiotic sex chromosome inactivation (MSCI), and piRNA production (Baudat *et al*, 2013; Li *et al*., 2013; Roeder & Bailis, 2000; Subramanian & Hochwagen, 2014). We show that pSpc is more vulnerable to NFYA-depletion compared to that at other stages of spermatogenesis. NFYA-depletion directly affects transcriptional programs regulating chromosome organization and DNA damage response in pSpc. In line with previous reports in other cell types (Benatti *et al*, 2011; Rigillo *et al*, 2021), NFYA-loss also leads to DNA damage increase, supporting a conserved role for NFYA in maintaining genome integrity. Intriguingly, although AMYB has been identified as a master regulator for both pachytene piRNA precursors and core piRNA biogenesis factors, including *Piwil1* (Li *et al*., 2013), our data suggest that NFYA may also contribute to piRNA biogenesis by activating piRNA biogenesis factors, including *Piwil1/2* (**Fig. 5I**), consistent with a previous report (Hou *et al*, 2012). Moreover, NFYA appears to act upstream of *Amyb* (**Fig. 2K**), although the effect on *Amyb* expression is modest. Together, these findings suggest that NFYA may act as an accessory regulator to support robust *Amyb* expression and piRNA production. However, the precise mechanism by which NFYA contributes to piRNA regulation remains to be elucidated in future studies.

Great progress has been made in identifying regulators of spermiogenesis, including RFX2 (Kistler *et al*, 2015) and SOX30 (Chen *et al*, 2018; Zhang *et al*, 2018). RFX2 knockout leads to defective cell adhesion and intracellular trafficking in Std (Kistler *et al*., 2015), whereas SOX30 knockout disrupts chromosome organization (Chen *et al*., 2018; Zhang *et al*., 2018). In contrast, we show that NFYA-depletion increases ROS levels and oxidative stress, triggering apoptosis in rStd, suggesting an essential role of NFYA in regulating antioxidative stress programs. Intriguingly, *Nfya*-KO in pancreatic β-cells has also been reported to induce oxidative stress and apoptosis (Liu *et al*, 2021), consistent with the mechanism revealed in this study.

Since NFYA is evolutionarily conserved across eukaryotes (Romier *et al*, 2002), and the core transcriptional programs during spermatogenesis, including piRNA production and MSCI, are also highly conserved (Brawand *et al*., 2011; Özata *et al*, 2019), it raises the possibility that NFYA’s role in spermatogenesis might be conserved in other species. Future work will determine whether this is indeed the case.

## Methods

### Animal

All experiments were conducted in accordance with the National Institutes of Health Guide for the Care and Use of Laboratory Animals and were approved by the Institutional Animal Care and Use Committee of Boston Children’s Hospital and Harvard Medical School (protocol no. IS00000270-6). All mice were maintained under specific pathogen-free conditions in a temperature-controlled (20–22 °C) and humidity-controlled (40–70%) environment with a 12-h light/dark cycle. NFYA-dTAG knock-in mice were generated as previously described (Yang *et al*., 2026). Genotyping was performed using mouse tails lysed in lysis buffer (50 mM Tris-HCl, 0.5% Triton and 400 μg/ml Proteinase K) at 55 °C overnight.

### *In vivo* microinjection of dTAG^V^-1

For *in vivo* protein degradation, dTAG^V^-1 was first dissolved in DMSO to generate a 40 mg/mL stock solution. For i.p. injection, the stock solution was diluted in sterile normal saline (NS) containing 10% Kolliphor EL (KEL) to a final concentration of 2 mg/mL. dTAG^V^-1 was administered at 40 mg/kg per mouse as a single dose. For the dTAG-IS approach, the stock solution was similarly diluted in vehicle (10% KEL in sterile NS) to the desired working concentration. Fast Green FCF (0.05% w/v) was added to the final working solution as a visual tracer to facilitate visualization of injection efficiency.

For the microinjection, adult male mice were anesthetized, and the testes were exteriorized through a midline abdominal incision. The efferent ducts were exposed by carefully retracting the epididymal adipose tissue. Approximately 10-15 μL of the dTAG^V^-1 solution was microinjected into the seminiferous tubules via the efferent duct using a Borosilicate glass micropipette with a fine tip. Following the injection, the efferent duct was ligated with an 8-0 suture at the segment proximal to the testis relative to the injection site. Successful filling of the tubular network was confirmed by the spread of the Fast Green dye throughout approximately two-thirds of the testis. Following the procedure, the testes and associated adnexal structures were carefully returned to the scrotal sac in the correct anatomical sequence. The abdominal wall was sutured, and the skin was closed with surgical clips to allow for recovery.

### Testis dissociation and digestion

Testes were harvested and decapsulated by removing the tunica albuginea to expose the seminiferous tubules. To isolate the tubules from interstitial cells, the tissue was initially incubated in HBSS containing 1 mg/mL Collagenase IV (Thermo, 17104019) at 34°C for 30 minutes with gentle agitation. The interstitial cell-depleted tubules were recovered using a 100 μm cell strainer, washed thoroughly with HBSS, and subsequently subjected to enzymatic dissociation in 0.1% Trypsin-EDTA (Thermo, 25200056) + 20 IU/ mL DNase I (NEB, M0303) at 4°C for 15 minutes. The digestion was neutralized by the addition of ice-cold FBS. Finally, the resulting cell suspension was passed through a 40 μm cell strainer to eliminate debris and ensure a high-quality single-cell suspension for downstream analysis.

### Fluorescence-activated cell sorting

For the isolation of Spc and rStd, cell suspensions were prepared in Cell Staining Buffer (BioLegend, 420201) supplemented with 0.5 mM EDTA. The cells were incubated with Hoechst 33342 (Thermo, H3570) at a final concentration of 5 μg/mL for 30 minutes at room temperature in the dark. For Spg identification, cells were stained with a cKIT antibody (eBioscience, 12-1171-83) at a 1:200 dilution. Following staining, cells were washed three times and resuspended in ice-cold PBS for sorting. Immediately prior to acquisition, Propidium Iodide (PI; Thermo, V13241) was added to the Spc and rStd samples at a 1:2000 dilution, while DAPI (BD, 564907) was added to the Spg samples at a 1:2000 dilution to assess cell viability. The gating strategy involved an initial selection based on forward and side scatter (FSC/SSC) to identify the target populations and exclude doublets. Dead cells were subsequently excluded based on PI or DAPI positivity. Spc and rStd populations were identified and gated according to their characteristic Hoechst blue/red fluorescence profiles, while Spg were identified based on cKIT expression levels. FACS were performed on a Sony SH800Z Cell Sorter. All flow cytometry data were processed and analyzed using FlowJo software.

### Spermatocyte chromosome spreading

Spermatocyte chromosome spreads were prepared using the hypotonic extraction buffer (HEB) method (Dia *et al*, 2017). Briefly, decapsulated seminiferous tubules were incubated for 20 minutes in ice-cold HEB (30 mM Tris-HCl, 50 mM Sucrose, 17 mM Citric Acid, 5 mM EDTA, 0.5 mM DTT; pH 8.2–8.4). The tubules were then minced in 100 mM sucrose to generate a cell suspension. Approximately 20 μL of this suspension was added to 100 μL of fixative (1% PFA, 0.15% Triton X-100; pH 9.2) on glass slides and spread evenly. Following overnight incubation in a humid chamber, slides were air-dried, washed sequentially with ddH_2_O and Photo-Flo solution, and followed by immunofluorescence (IF) staining steps or stored at −80°C.

### DCFH-DA assay

Intracellular ROS levels were determined using the Cellular ROS Assay Kit (Abcam, ab113851) following the manufacturer’s protocol. Germ cells were harvested to ensure a single-cell suspension and first stained with Hoechst 33342 following the protocol in “Fluorescence-activated cell sorting”. 20 mM DCFH-DA stock solution was diluted in PBS to a working concentration of 20 μM. Cells were then stained with the diluted DCFH-DA solution for 30 minutes at 34°C in the dark. Cells were washed three times before flow cytometry analysis.

### Immunofluorescence (IF)

For IF staining, the slides were first blocked and permeabilized with 5% Bovine Serum Albumin (BSA) and 0.5% Triton X-100 in PBS for 30 min at room temperature. The slides were then incubated overnight at 4°C with a primary antibody. Secondary antibodies including Alexa Fluor 647 (Thermo Scientific, A-21447, 1:500), donkey anti-rabbit IgG (H + L) secondary antibody, Alexa Fluor 488 (Thermo Scientific, A-21206, 1:500) and Donkey anti-mouse IgG Secondary Antibody, Alexa Fluor 568 (Fisher Scientific, A10037, 1:500) were used in this study. After washing three times with PBS, the slides were incubated with fluorophore-conjugated secondary antibodies for 1 hour at room temperature. Nuclei were counterstained with Hoechst 33342. Acrosome was stained with 1 μg/ml PNA (Biotium, 29061). Confocal imaging was performed with Zeiss LSM800.

### TUNEL assay

Apoptotic cells were detected using the One-step TUNEL Apoptosis Kit (Green, AF488) (Elabscience, E-CK-A321) according to the manufacturer’s instructions. Briefly, the slides were permeabilized with Proteinase K. The slides were then washed and incubated with TdT Equilibration Buffer at 37°C for 20 minutes and incubated with the TUNEL reaction mixture for 60 minutes at 37°C in a humidified dark chamber. After washing with PBS, nuclei were counterstained with Hoechst 33342.

### CUT&RUN and RNA-seq library preparation and sequencing

CUT&RUN was conducted on male germ cells according to established protocols with optimizations (Zhou *et al*., 2025). Briefly, approximately 5×10^4^ cells were harvested and resuspended in 50 μL of washing buffer (20 mM HEPES pH 7.5, 150 mM NaCl, 0.5 mM spermidine, and 1x protease inhibitor). The cells were then coupled with pre-activated Concanavalin A-conjugated magnetic beads (Polysciences, 86057-3) for 10 minutes at room temperature. Following bead binding, samples were incubated overnight at 4°C with a primary antibody against HA-Tag (1:100, Abcam, ab9110). After primary antibody incubation, the cell-bead complexes were treated with 2.8 ng/μL of home-made pA-MNase for 2h at 4°C. Targeted chromatin cleavage was initiated by adding 200 μL of ice-cold 0.5 μM CaCl_2_ and maintained at 4°C for 20 minutes. The reaction was stopped with the addition of 23 μL of 10x stop buffer (1.7 M NaCl, 20 mM EGTA, 100 mM EDTA, 0.02% Digitonin, 250 μg/mL glycogen, and 250 μg/mL RNase A). Digested DNA fragments were released into the supernatant by incubation at 37°C for 15 minutes. Subsequently, the samples underwent protein digestion with 10% SDS and 20 mg/mL Proteinase K at 55°C for at least 1 hour. Purified DNA was obtained via phenol-chloroform extraction followed by ethanol precipitation. Sequencing libraries were prepared with the NEBNext Ultra II DNA library preparation kit for Illumina (New England Biolabs, E7645S). RNA-seq library was prepared by SMART-Seq® Stranded Kit (Takara Bio, 63444) using 5×10^4^ cells. All libraries were sequenced by NextSeq 550 system (Illumina) with paired-end 75-bp reads (Dataset EV6).

### RNA-seq data processing

Raw sequencing reads were trimmed using Trimmomatic (Bolger *et al*, 2014) (v0.39) to remove sequencing adaptors, and subsequently mapped to the GRCm38 genome using STAR (Dobin *et al*, 2012) (v2.7.8a). Gene expression was quantified with featureCounts (Liao *et al*, 2013) (v2.0.1) by counting reads mapped to each gene. Then, edgeR (Robinson *et al*, 2009) (v3.32.1) was employed for normalization and differential expression analysis. Genes with RPKM lower than 1 were defined as lowly expressed genes and excluded from the differential expression analysis. Differentially expressed genes (DEGs) were identified using likelihood ratio test (glmFit and glmLRT functions from edgeR). Genes with a false discovery rate (FDR) below 0.05 and an absolute value of fold change greater than 2 were defined as DEGs. DEGs identified from diff-Spg, pSpc, and rStd were used for downstream stage-specific analyses and visualization (Supplemental Tables S1-S2 and S4). GO enrichment was performed using the R package clusterProfiler (Yu *et al*, 2012). GSEA was performed using GSEA software (v4.0.3) (Mootha *et al*, 2003; Subramanian *et al*, 2005).

### CUT&RUN data analysis

Raw sequencing reads were trimmed using Trimmomatic (Bolger *et al*., 2014) (v0.39) to remove sequencing adaptors, and subsequently aligned to the GRCm38 reference genome using bowtie2 (Langmead & Salzberg, 2012) (v2.4.2) with parameters: --local --very-sensitive-local -- no-unal --no-mixed --no-discordant --dovetail -I 10 -X 700 --soft-clipped-unmapped-tlen. PCR duplicates were removed by Picard MarkDuplicates (Broad Institute, 2019) (v2.23.4) and reads with a mapping quality below 30 were removed. The mapped reads were further filtered to only retain proper paired reads with fragment length between 10 and 120 bp. Then, MACS2 (Zhang *et al*, 2008) (v2.2.7.1) was used to call significant peaks with parameters “-f BAMPE -B --SPMR - q 0.05 -g mm --keep-dup all”. To obtain highly conserved NFYA binding sites, only peaks present in both biological replicates were defined as binding sites. The signal tracks were generated with deeptools (Ramírez *et al*, 2016) bamCoverage (v3.5.1) with bin size of 1 and normalized using CPM. Peak annotation was performed using the ChIPseeker (Yu *et al*, 2015) R package. Peaks located within ±2,500 bp of transcription start sites (TSS) were classified as promoter peaks, whereas peaks outside this region were considered distal peaks. The heatmaps of binding profiles were calculated with deeptools (Ramírez *et al*., 2016) computeMatrix (v3.5.1) using bigwig signal tracks as input and bin size of 10 and visualized in R with profileplyr and EnrichedHeatmap (Gu *et al*, 2018) packages.

### Transcription factor enrichment analysis

Transcription factor enrichment analysis was performed using the ChIP-Atlas “Enrichment” function (https://chip-atlas.org) (Oki *et al*, 2018; Zou *et al*, 2022; Zou *et al*, 2024). Genomic regions of interest were submitted as BED files and compared against a comprehensive collection of publicly available ChIP-seq peak datasets. Enrichment of transcription factor binding was evaluated based on the overlap between input regions and ChIP-seq peaks from each dataset. Statistical significance was assessed using Fisher’s exact test, and *p* values were adjusted for multiple comparisons using the Benjamini-Hochberg method. Transcription factors with adjusted *p* value less than 0.05 were considered significantly enriched.

### Motif occurrence analysis

Motif occurrence analysis was performed using HOMER (Heinz *et al*, 2010) (v4.11) annotatePeaks.pl with parameters mm10 -size −500,500 -hist 20 -ghist for regions around gene TSSs. Motif files were downloaded from the JASPAR database (Castro-Mondragon *et al*, 2022) and manually converted to HOMER motif format. The JASPAR (Fornes *et al*, 2020) motif ID for NFYA analyzed in this study is MA0060.1, and a log odds detection threshold of 6.0 was applied. The motif occurrence matrix was visualized in R with the EnrichedHeatmap (Gu *et al*., 2018) package.

### Prediction of NFYA-mediated promoter-enhancer interactions

To predict NFYA-mediated promoter-enhancer interactions, pSpc RNA-seq and NFYA CUT&RUN, as well as public datasets of the pachytene Spc, including ATAC-seq (Maezawa *et al*., 2018) and H3K27ac ChIP-seq (Chen *et al*, 2020), were collected as the input for Activity-by-Contact (ABC) model (v0.2) (Fulco *et al*., 2019). Enhancers within 5 Mb and with scores ≥0.02 were assigned to target genes (Dataset EV6).

## Quantification and statistical analysis

Student’s *t* tests or multiple *t* tests for graph analysis were performed with GraphPad Prism (v8.3.0). Data distribution was assumed to be normal, but this was not formally tested. For the IF experiments, at least three independent repetitions were performed with consistent results, and representative data were presented. *p* values < 0.05 were considered statistically significant. No statistical methods were used to pre-determine sample sizes, but our sample sizes are similar to or greater than those reported in previous publications. No data was excluded from the analysis.

## Resource availability

Further information and requests for resources and reagents should be directed to the lead contact, Yi Zhang.

## Data availability

Sequencing data that support the findings of this study have been deposited in the Gene Expression Omnibus (GEO) under accession code GSE326586 (reviewer token: cnaxmukirditnsf). Public data, including RNA-seq (Horisawa-Takada *et al*., 2021; Kojima *et al*., 2019; Lin *et al*., 2025), ATAC-seq (Maezawa *et al*., 2018), H3K27ac ChIP-seq (Chen *et al*., 2020) and CUT&RUN (Cecchini *et al*, 2023; Kojima *et al*., 2019; Xu *et al*., 2022), were used in this study. The details are listed in Dataset EV7. Source data are provided with this paper. This paper does not report original code.

## Acknowledgments

We thank members of the Zhang lab for discussion during the study; Chengjie Zhou and Yota Hagihara for commenting on the manuscript. This project was supported by the HHMI. YZ is an investigator of the Howard Hughes Medical Institute.

## Author contributions

Boyan Wang: Methodology; Investigation; Validation; Visualization; Writing—original draft; Writing—review and editing. Shan Jiang: Formal analysis; Software; Data curation; Visualization; Writing—original draft; Writing—review and editing. Qianying Yang: Resources; Investigation; Visualization; Writing—original draft; Writing—review and editing. Yi Zhang: Conceptualization; Funding acquisition; Project administration; Supervision; Writing—original draft; Writing—review and editing.

