## Supplemental_Figures for "NFYA regulates spermatogenesis through multifaceted chromatin interactions"

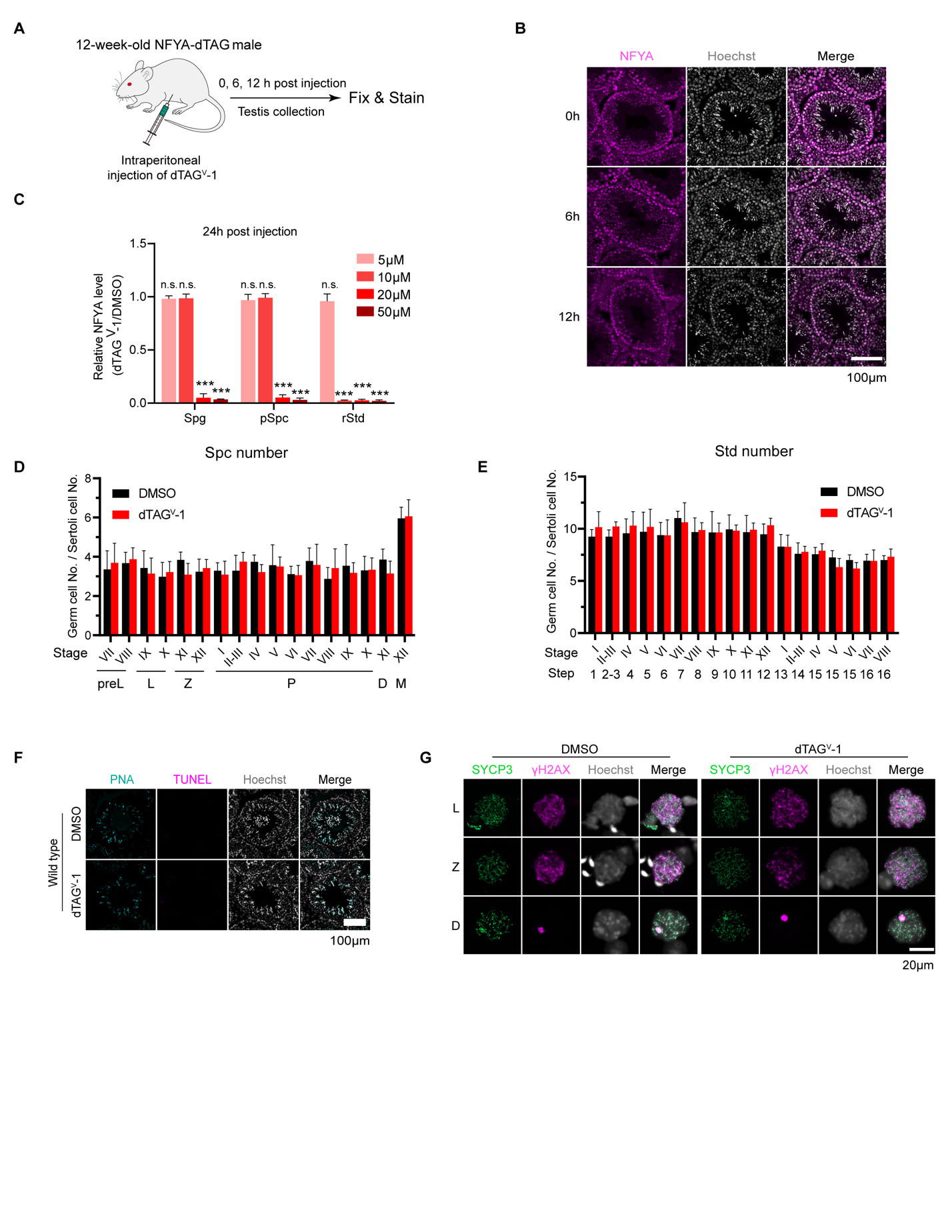

**Figure EV1. Establishment of dTAG-induced rapid NFYA degradation system in testis.**

(A) Experimental workflow for acute NFYA protein degradation *in vivo* via intraperitoneal (i.p.) injection of dTAG^V^-1. (B) IF staining of HA-tagged NFYA (magenta) in NFYA-dTAG testis sections collected at different time points after dTAG^V^-1 administration. Scale bar, 100 μm. (C) Quantification of NFYA mean fluorescence intensity relative to DMSO control in spermatogonia (Spg), pachytene spermatocytes (pSpc), and round spermatids (rStd) after treatment with different concentrations of dTAG^V^-1. (D) Quantification of Spc number per Sertoli cell across seminiferous epithelial stages in DMSO and dTAG^V^-1 treated testes. L, leptotene; Z, zygotene; P, pachytene; D, diplotene; M, meiotic division. (E) Quantification of spermatid number per Sertoli cell across round spermatid (rStd) and elongating spermatid (eStd) steps in DMSO and dTAG^V^-1 treated testes. (F) TUNEL assay (magenta) on testis sections from DMSO and dTAG^V^-1 treated mice. Scale bar, 100 μm. (G) IF staining of SYCP3 (green), γH2AX (magenta) in Spc spreads from DMSO or dTAG^V^-1 treated testes at leptotene (L), zygotene (Z), and diplotene (D) stages. Scale bar, 20 μm. Quantitative data are shown as mean ± SEM; ****p* < 0.001, multiple *t* tests.

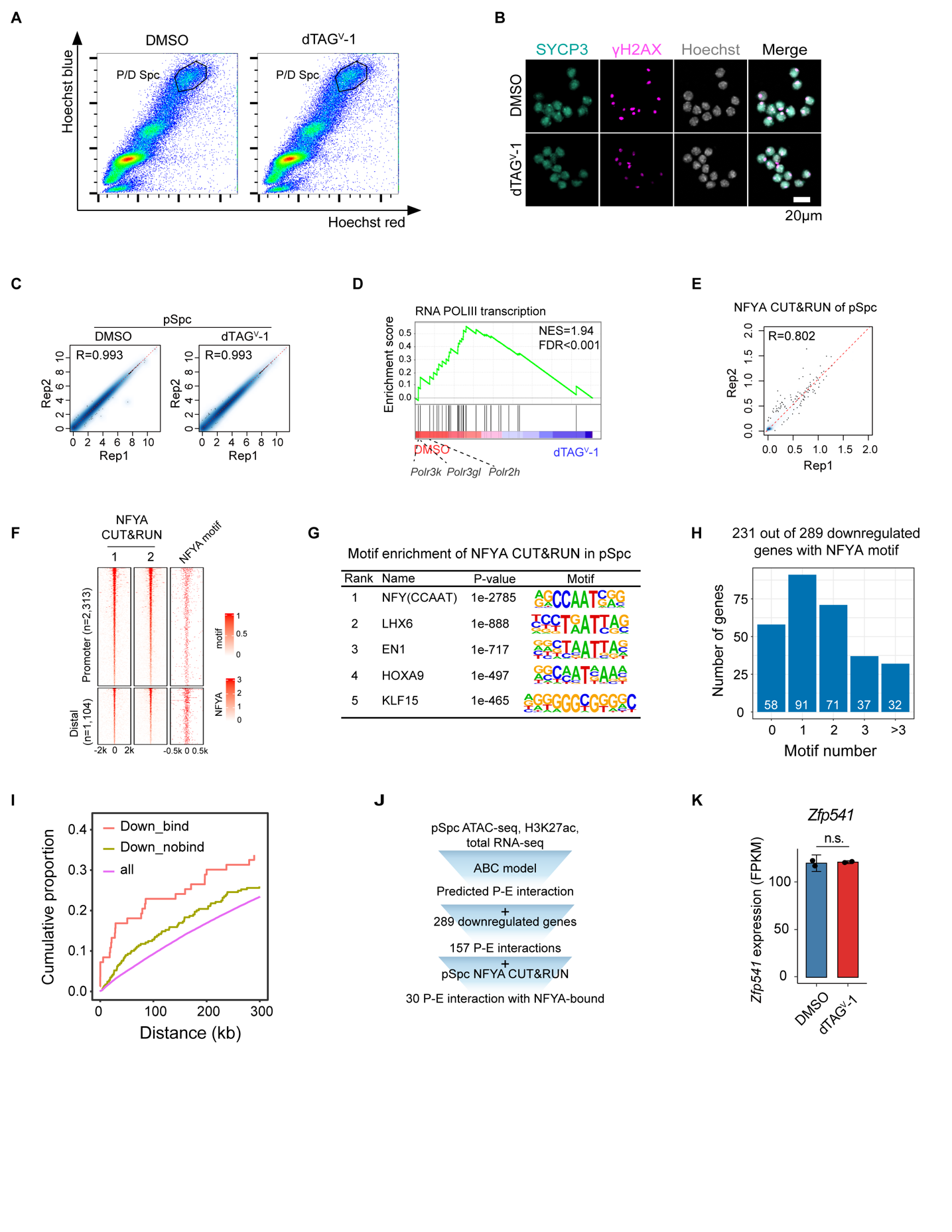

**Figure EV2. NFYA directly regulates chromosome organization and DNA damage response programs in pachytene spermatocytes.**

(A) Representative FACS plots of seminiferous tubule cells showing Hoechst Blue and Hoechst Red staining profiles in the DMSO and dTAG^V^-1 treated groups. The gated population corresponds to pachytene/diplotene (P/D) Spc. (B) Representative IF images of pachytene spermatocytes from the DMSO and dTAG^V^-1 treated groups stained for SYCP3 (cyan), γH2AX (magenta), and Hoechst 33342 (gray). Scale bar, 20 μm. (C) Correlation between biological replicates of total RNA-seq in pSpc. The x- and y-axes show Log (CPM+1) values. (D) GSEA showing significant downregulation of genes associated with RNA polymerase III transcription in dTAG^V^-1 treated pSpc. (E) Correlation between biological replicates of NFYA CUT&RUN in pSpc. Each dot represents a 5 kb bin. (F) Heatmaps showing NFYA CUT&RUN signals at promoter and distal NFYA-binding regions in pSpc, together with NFYA motif occurrence. (G) Top five enriched transcription factor binding motifs in NFYA-bound regions in pSpc. Based on *p* values, the NFYA motif is much more strongly enriched than the other motifs. (H) The numbers of NFYA motifs identified in the promoters of 289 downregulated genes in pSpc. (I) Cumulative distribution of the distance from gene TSSs to the nearest distal NFYA peaks in downregulated NFYA-bound genes, downregulated NFYA-unbound genes, and all genes as background. (J) Schematic workflow for prediction of NFYA-mediated promoter-enhancer interactions. pSpc ATAC-seq, H3K27ac, and total RNA-seq data were integrated using the ABC model, and the predicted interactions were intersected with the 289 downregulated genes and pSpc NFYA CUT&RUN data, identifying 30 promoter-enhancer interactions with direct NFYA binding. (K) Bar plot showing *Zfp541* expression levels in DMSO and dTAG^V^-1 treated pSpc. n.s., FDR > 0.05.

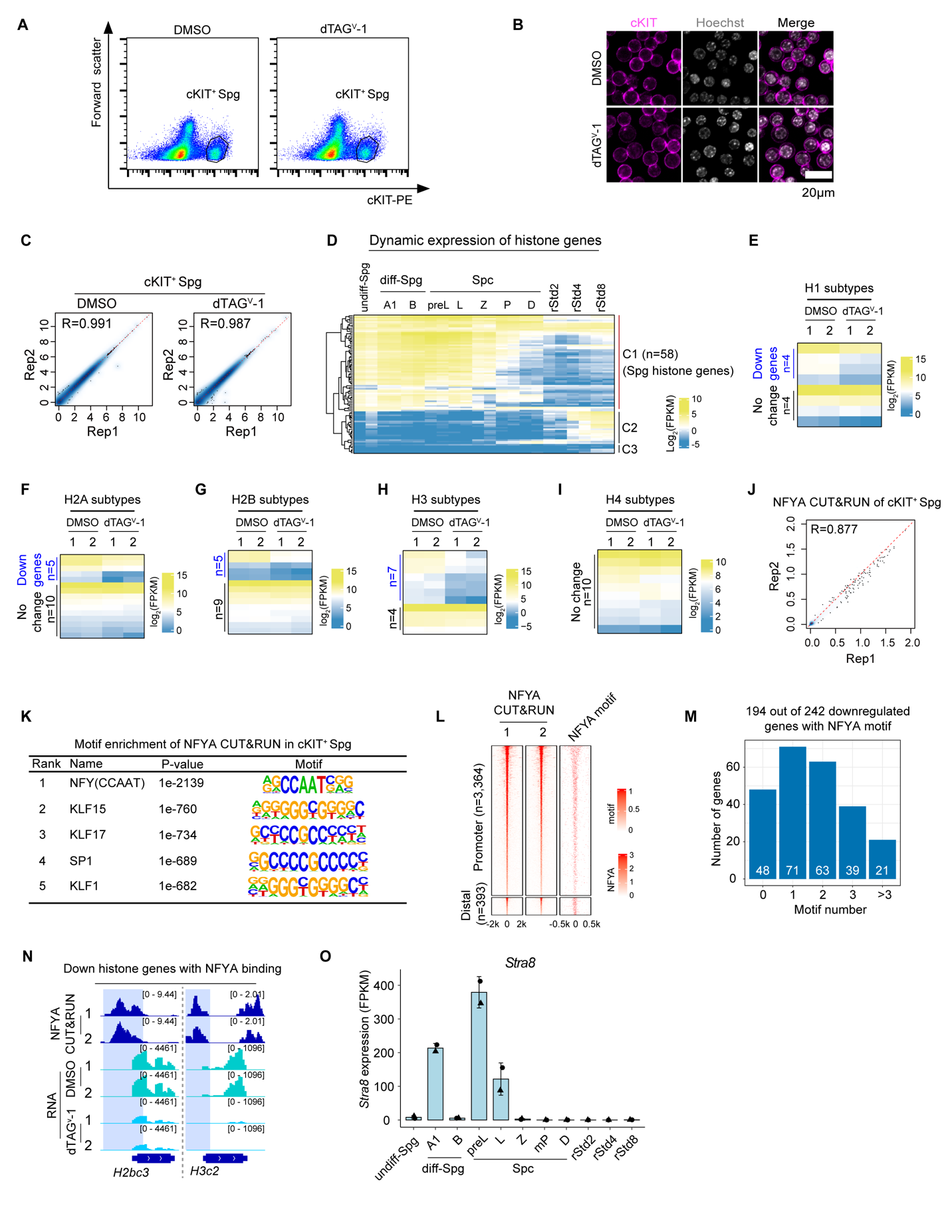

**Figure EV3. NFYA directly regulates cell cycle progression and histone gene upregulation in differentiating spermatogonia.**

(A) Representative FACS plots of seminiferous tubule cells showing forward scatter versus cKIT-PE staining profiles in the DMSO and dTAG^V^-1 treated groups. The gated population represents cKIT^+^ Spg. (B) Representative IF images of cKIT^+^ Spg from the DMSO and dTAG^V^-1 treated groups stained for cKIT (magenta) and Hoechst 33342 (gray). Scale bar, 20 μm. (C) Correlation analysis of total RNA-seq between biological replicates of cKIT^+^ Spg. The x- and y-axes show Log (CPM+1) values. (D) Heatmap showing the dynamic expression patterns of histone genes across undiff-Spg, diff-Spg, Spc, and rStd. Histone genes enriched in Spg are highlighted (C1, n=58). (E-I) Heatmaps showing expression changes of histone H1 (E), H2A (F), H2B (G), H3 (H), and H4 (I) genes in cKIT^+^ Spg after NFYA depletion. (J) Correlation analysis between biological replicates of NFYA CUT&RUN in cKIT^+^ Spg. Each dot represents a 5 kb bin. (K) Top five transcription factor binding motifs enriched in NFYA-bound regions in cKIT^+^ Spg. The NFYA motif showed the strongest enrichment based on *p* value. (L) Heatmaps showing NFYA CUT&RUN signals at promoter-proximal and distal NFYA-binding regions in cKIT^+^ Spg, together with NFYA motif occurrence. (M) Distribution of NFYA motif numbers in the promoters of the 242 downregulated genes in cKIT^+^ Spg. (N) Genome browser views of representative histone genes with direct NFYA binding that are downregulated upon NFYA depletion, showing NFYA CUT&RUN and RNA-seq signals in cKIT^+^ Spg treated with DMSO or dTAG^V^-1. (O) Bar plot showing *Stra8* expression levels across spermatogenesis stages based on RNA-seq data from GSE242515.

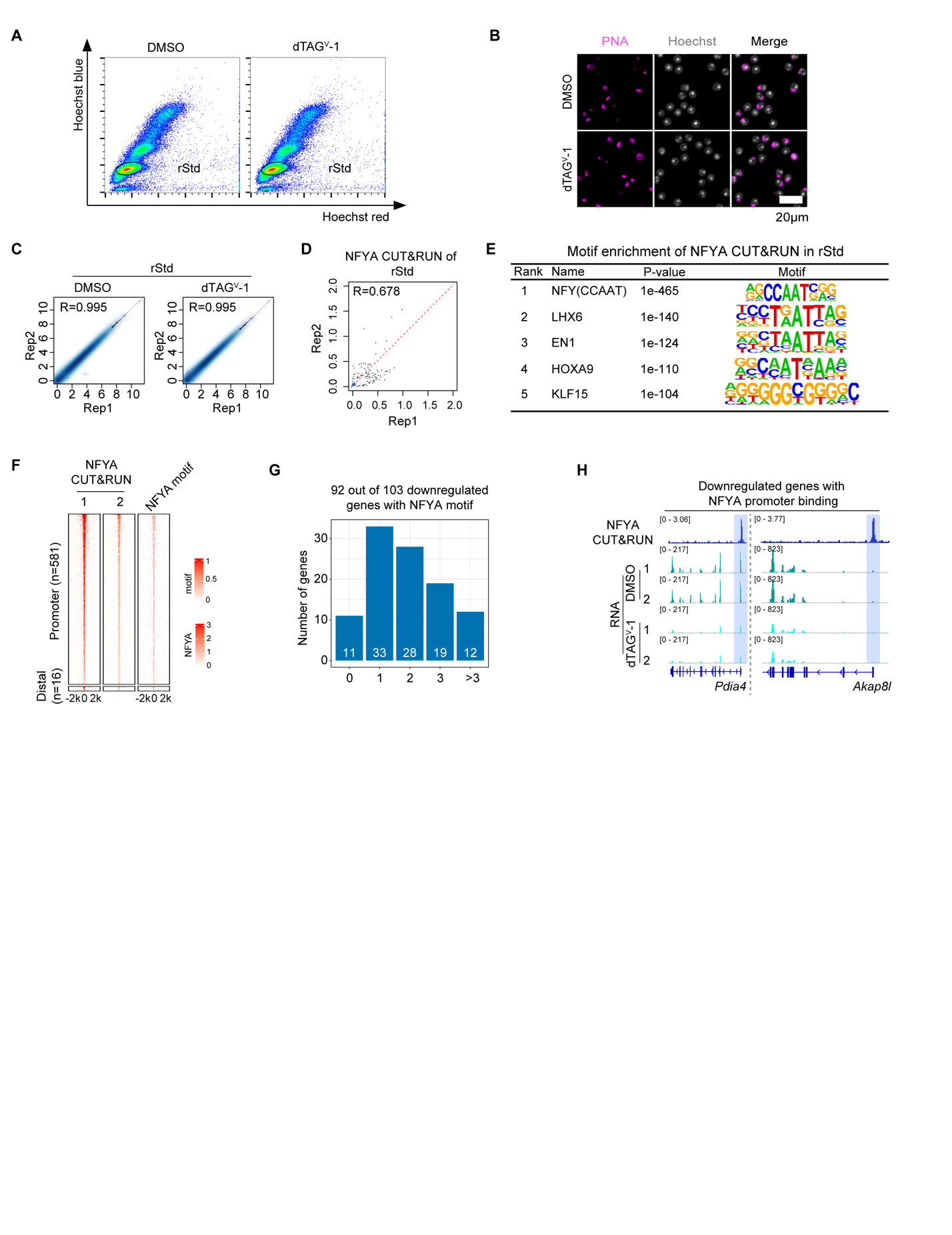

**Figure EV4. NFYA directly regulates oxidative stress response genes in round spermatids.**

(A) Representative FACS plots of seminiferous tubule cells showing Hoechst Blue versus Hoechst Red staining profiles in the DMSO and dTAG^V^-1 treated groups. The gated population represents round spermatids (rStd). (B) Representative fluorescence images of round spermatids from the DMSO and dTAG^V^-1 treated groups stained with PNA (magenta) and Hoechst 33342 (gray). Scale bar, 20 μm. (C) Correlation analysis between biological replicates of total RNA-seq in rStd. The x- and y-axes show Log (CPM+1) values. (D) Correlation analysis between biological replicates of NFYA CUT&RUN in rStd. Each dot represents a 5 kb bin. (E) Top five transcription factor motifs enriched in NFYA-bound regions in rStd. The NFYA motif showed the highest enrichment based on *p* value. (F) Heatmaps showing NFYA CUT&RUN signals at promoter-proximal and distal NFYA-binding regions in rStd, together with NFYA motif occurrence. (G) Distribution of NFYA motif numbers in the promoters of the 103 downregulated genes in rStd. (H) Genome browser views of representative downregulated genes with direct NFYA promoter binding, showing NFYA CUT&RUN and RNA-seq signals in rStd treated with DMSO or dTAG^V^-1.

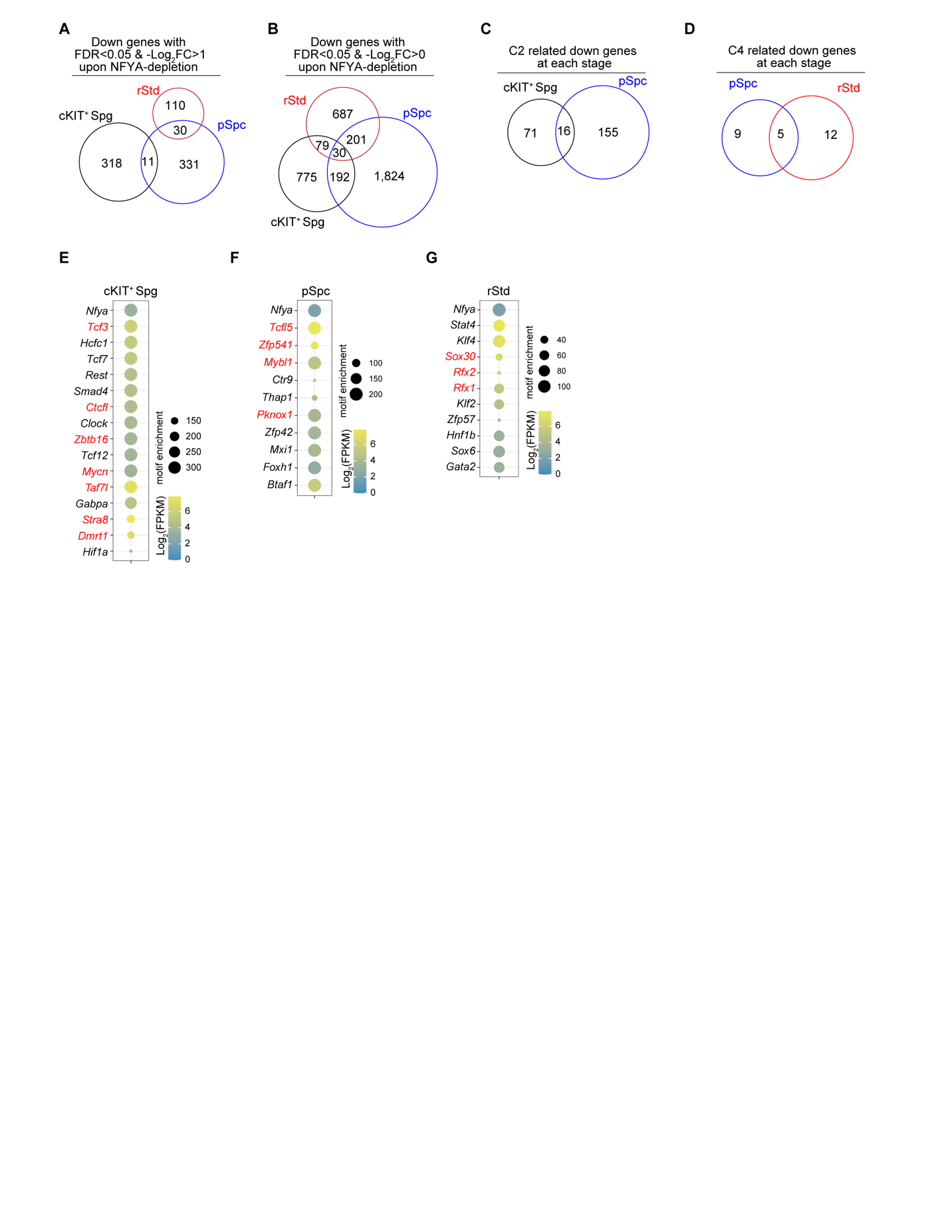

**Figure EV5. Stage-specific NFYA-regulated transcriptional programs and motif features across spermatogenesis.**

(A) Venn diagram showing the overlap of downregulated genes across cKIT^+^ Spg, pSpc, and rStd after NFYA depletion, using the threshold FDR<0.05 and -Log_2_FC>1. (B) Venn diagram showing the overlap of downregulated genes across cKIT^+^ Spg, pSpc, and rStd after NFYA depletion, using a more permissive threshold FDR<0.05 and -Log_2_FC>0. (C-D) Venn diagrams showing the overlap of cluster-associated NFYA-binding-related downregulated genes across cKIT^+^ Spg, pSpc, and rStd for NFYA-binding clusters C2 and C4. (E-G) Motif enrichment analysis of functional NFYA binding regions in cKIT^+^ Spg, pSpc, and rStd.
